# Selective attention is insensitive to reward and to dopamine in Parkinson’s disease

**DOI:** 10.1101/2021.04.22.440852

**Authors:** Matthew Pilgrim, Zhen-Yi Andy Ou, Madeleine Sharp

## Abstract

Patients with Parkinson’s disease exhibit reduced reward sensitivity in addition to early cognitive deficits, among which attention impairments are common. Attention allocation is controlled at multiple levels and recent work has shown that reward, in addition to its role in the top-down goal-directed control of attention, also guides the automatic allocation of attention resources, a process thought to rely on striatal dopamine. Whether Parkinson’s patients, due to their striatal dopamine loss, suffer from an inability to use reward information to guide the allocation of their attention is unknown. To address this question, we tested Parkinson’s patients (n=43) ON and OFF their dopaminergic medication, and compared them to a group of older controls (n=31). We used a standard two-phase attention capture task in which subjects were first implicitly trained to make colour-reward associations. In the second phase, the previously reward-associated colours were used as distractors in a visual search task. We found that patients did not use reward information to modulate their attention; they were similarly distracted by the presence of low and high-reward distractors. However, contrary to our predictions, we did not find evidence that dopamine modulated this inability to use reward to guide attention allocation. Additionally, we found slightly increased overall distractibility in Parkinson’s patients compared to older controls, but interestingly, the degree of distractibility was not influenced by dopamine replacement. Our results suggest that loss of reward-guided attention allocation may contribute to early attention deficits and raise the possibility that this inability to prioritize cognitive resource allocation could contribute to executive deficits more broadly in Parkinson’s disease.

## Introduction

Where we focus our attention plays a critical role in downstream cognitive processes. The ability to automatically and selectively attend to some environmental stimuli over others allows us to filter incoming information. This necessarily influences what enters memory, what we learn about, and what we are subsequently drawn to in our decisions (Johnston & Dark, 1986). Given our limited cognitive resources, the ability to selectively attend to information that is behaviourally relevant is a key step towards aligning cognitive resource allocation with our goals (Franconeri et al., 2013; Scalf et al., 2013). Mild attention deficits are well-established in early Parkinson’s disease but whether these reflect an inability to automatically prioritize the allocation of attentional resources to what matters most is unknown (Dirnberger & Jahanshahi, 2013; Kudlicka et al., 2011). Meanwhile, early Parkinson’s disease, a stage when cognitive function remains relative intact, is a time when subtle failures of selective attention allocation could potentially have an important impact on behaviour more broadly, by influencing downstream cognitive processes and their content.

Much of the work on early attention deficits in Parkinson’s patients has focused on overall attentional capacity (Aarsland et al., 2010; Dirnberger & Jahanshahi, 2013; Weintraub et al., 2015) rather than the content or the focus of attention. One important exception is the body of work on attention set-shifting in Parkinson’s patients (Fallon et al., 2013; Fallon et al., 2016; Owen et al., 1993; Slabosz et al., 2006), which has examined the use of top-down control mechanisms to voluntarily guide attention allocation towards rewarding stimuli (Corbetta & Shulman, 2002; Gazzaley & Nobre, 2012). These studies have demonstrated that Parkinson’s patients are impaired at using top-down control strategies to guide their attention towards reward, and that they are especially impaired when they are required to shift the focus of their attention because of changing reward contingencies. However, these studies do not address the more rapid and automatic process by which some elements of the environment are selected for attentional processing over others.

Meanwhile, recent evidence has highlighted an important role for reward in guiding the automatic selectivity of attention, a process which is thought to depend on striatal dopamine (Anderson et al., 2011; Anderson et al., 2016, 2017). This bears particular relevance to Parkinson’s disease where dopamine-dependent reward processing is disrupted (Bódi et al., 2009; Frank et al., 2004; Maril et al., 2013; Palminteri et al., 2009; Rutledge et al., 2009; Skvortsova et al., 2017). One approach to demonstrating the way in which reward can modulate the focus of attention has been to associate reward with stimuli that are not relevant to the task at hand, and instead function as distractors. Across a number of studies, it has been shown that the magnitude of reward associated with a distractor can enhance the degree to which it causes distraction, such that distractors associated with high reward are more distracting that those associated with low reward (Anderson et al., 2011; Anderson, 2013; Anderson & Halpern, 2017; Anderson & Yantis, 2013). These findings support the notion that reward signals in the environment can orient attention towards stimuli in a manner that is distinct from top-down or bottom-up control. Studies looking to uncover the neural mechanisms of this process have indicated that the striatum, and more specifically, dopaminergic pathways, are involved (Anderson, 2017; Anderson et al., 2014, 2016, 2017), and that reward-related activity in dopaminergic pathways is associated with enhanced representation of stimuli in sensory cortices (Hickey & Peelen, 2015).

It is well established, across a number of different paradigms, that Parkinson’s patients have reduced reward sensitivity related to dopamine loss. Much of this work has focused on how this manifests as reduced learning from reward (Bódi et al., 2009; Cools et al., 2006; Frank et al., 2004; Palminteri et al., 2009; Rutledge et al., 2009; Skvortsova et al., 2017). However, even reward sensitivity measured in isolation of any dependent cognitive process is reduced in Parkinson’s patients (Manohar & Husain, 2015; Muhammed et al., 2016).This raises the possibility that altered reward signaling could have consequences on cognitive processing *beyond* its effect on learning and motivation. Indeed, there is evidence that reward helps guide the automatic prioritization of cognitive resource allocation across a number of domains, including working memory (Gong & Li, 2014; Infanti et al., 2015; Klink et al., 2017) and episodic memory (Adcock et al., 2006; Braun et al., 2018), and that this reward-guided prioritization process is lost in Parkinson’s patients (Sharp et al., 2020). Whether the reward-guided allocation of attention resources is similarly impaired in Parkinson’s disease in a dopamine-dependent manner is not known.

To address this question, we used a standard two-stage attention capture task, which has been previously used to show the role of reward in guiding attention (Anderson et al., 2011; Anderson, 2017; Anderson et al., 2016, 2017; Anderson & Halpern, 2017; Anderson & Yantis, 2013). The first phase is a reward-association paradigm where different stimuli are paired with either a low or a high reward. The second phase is an attention test where the stimuli previously associated with reward now act as goal-irrelevant distractors to draw attention away from targets. To assess the role of dopamine, we tested patients with Parkinson’s disease in a within-subject design, both ON and OFF dopaminergic medication, and compared them to older controls. As expected, we found that Parkinson’s patients did not use reward information to allocate their attention; patients were distracted by both the high-reward and the low-reward distractors to a similar degree. Contrary to our predictions, however, we did not find that dopamine modulated the effect of reward on attention, nor the overall distractibility; patients both ON and OFF were insensitive to the level of reward associated with the distractors. These findings suggest that loss of reward-guided attention allocation may contribute to early attention deficits in Parkinson’s disease and suggest that dopamine may not uniformly play a role across cognitive processes in the prioritization of resource allocation.

## Methods

### Patients and control subjects

Forty-three Parkinson’s disease patients (13 females, mean ± SD age = 63.8 ± 6.4) and 31 control subjects (21 females, mean ± SD age = 63.8 ± 7.9) were recruited to participate in our study. Patients were recruited from the Movement Disorders Clinic at the Montreal Neurological Institute, community groups and the Parkinson Quebec Network, a registry of Parkinson’s patients interested in research who have been referred by movement disorder specialists. Control subjects were recruited from spouses and friends of patients, community groups and social media posts. Controls had no major health issues or current neurological diagnoses. All subjects gave informed written consent and were compensated for their participation. Disease duration ranged from 0.42 −14.25 years (Mean years = 4.75 ± 3.25). All patients were taking levodopa, 6 patients were additionally taking a dopamine agonist (either pramipexole or rotigotine). See Table 1 for detailed demographic and clinical information. Comparing demographics across groups with Welch-approximated two-sample T-tests (Welch, 1947) and Chi-squared tests, we note that patients had fewer years of education than controls (p = 0.031) and that there were fewer women in the Parkinson’s group than in the control group (p = 0.003). To control for these differences, we included sex and education as covariates in our analysis. The study was approved by the McGill University Health Centre Research Ethics Board.

**Table 1.** Demographic and neuropsychological information. MoCA = Montreal Neurological Assessment, Verbal Fluency is taken from the Language section of the MoCA. Values presented are mean (SD). * p<0.05, ** p<0.01.

| Measure | Parkinson's patients (N = 43) | Controls (N = 31) | P-Value |
| --- | --- | --- | --- |
| Age | 63.8 (6.4) | 63.8 (7.9) | .996 |
| Education, years | 15.2 (3.5) | 17.1 (2.7) | .009 ** |
| **Disease duration, years** | 4.8 (3.3) | NA | NA |
| **Total PD Medication, mg** | 622.3 (363.3) | NA | NA |
| **Percent Female** | 30% | 68% | .003 ** |
| **MoCA** | 26.7 (2.6) | 27.8 (1.5) | .031 * |
| **Verbal Fluency (MoCA)** | 12.4 (4.2) | 13.8 (3.4) | .100 |
| **Digit Span Test** | 11.2 (2.3) | 12.1 (2.0) | .077 |
| **Symbol Digit Modalities Test** | 40.1 (10.7) | 47.4 (8.5) | .002 ** |
| **Geriatric Depression Scale** | 8.6 (6.1) | 5.4 (5.4) | .114 |
| **Apathy Evaluation Scale** | 58.4 (8.2) | 58.9 (12.2) | .825 |

### General procedure and medication manipulation

All subjects came to the lab for two sessions and, to minimize practice effects, the interval between sessions was at least one and a half months. At both sessions, subjects completed the full neuropsychological battery (described below) and a behavioral task which was divided into two phases: the reward association phase and the attention test phase (described in detail below). All sessions were conducted in the morning, starting between 9 and 10 AM to allow us to more easily control the timing of medications and to control for circadian factors. For Parkinson’s disease patients, the OFF medication session was conducted after an overnight withdrawal (minimum 15 hours) from dopamine medications, and the ON session was conducted with patients having taken their medication one hour prior to the start of testing. The order of these sessions was counterbalanced across subjects. Fifteen Parkinson’s patients did not complete their second session: eight missed their OFF session and seven missed their ON session. Three older controls did not complete their second session. All of these subjects were still included in the analysis such that we have 28 patients and 28 controls with both sessions, and 15 patients and 3 controls with only one session. See **Supplementary Table 1** for demographic comparisons between the ON and OFF samples.

### Neuropsychological battery

All subjects were administered a neuropsychological battery to establish baseline cognitive functioning (**Table 1**). This battery included the Montreal Cognitive Assessment (MoCA) (Nasreddine et al., 2005), the Digit Span (Lezak et al., 2004) and the Symbols Digit Modalities Test (SDMT) (Smith, 1973). Subjects were also administered the Geriatric Depression Scale (Yesavage, 1988) and the Apathy Evaluation Scale (Marin et al., 1991). Patients and controls were compared on their various neuropsychological scores using Welch-approximated two-sample T-tests (Welch, 1947). Patients scored 1.1 points lower on the MoCA (p = 0.031) and 7.3 points lower on the SDMT (p = 0.002). The SDMT has been used in studies of other neurodegenerative patient populations as an overall marker of cognitive ability that is sensitive to decline (Lemiere et al., 2004; Rodrigues et al., 2009). For this reason, to account for differences in cognitive ability we chose to include performance on the SDMT in our models.

### Task

We made minor adjustments to a task commonly used to measure the influence of reward on attention (Anderson et al., 2011; Anderson, 2013; Anderson et al., 2014, 2016, 2017; Anderson & Halpern, 2017; Anderson & Yantis, 2013) in order to make it more suitable for an older population. At each session, subjects performed a task with two phases: the reward association phase and the attention test phase (**Figure 1**).

**Figure 1.**
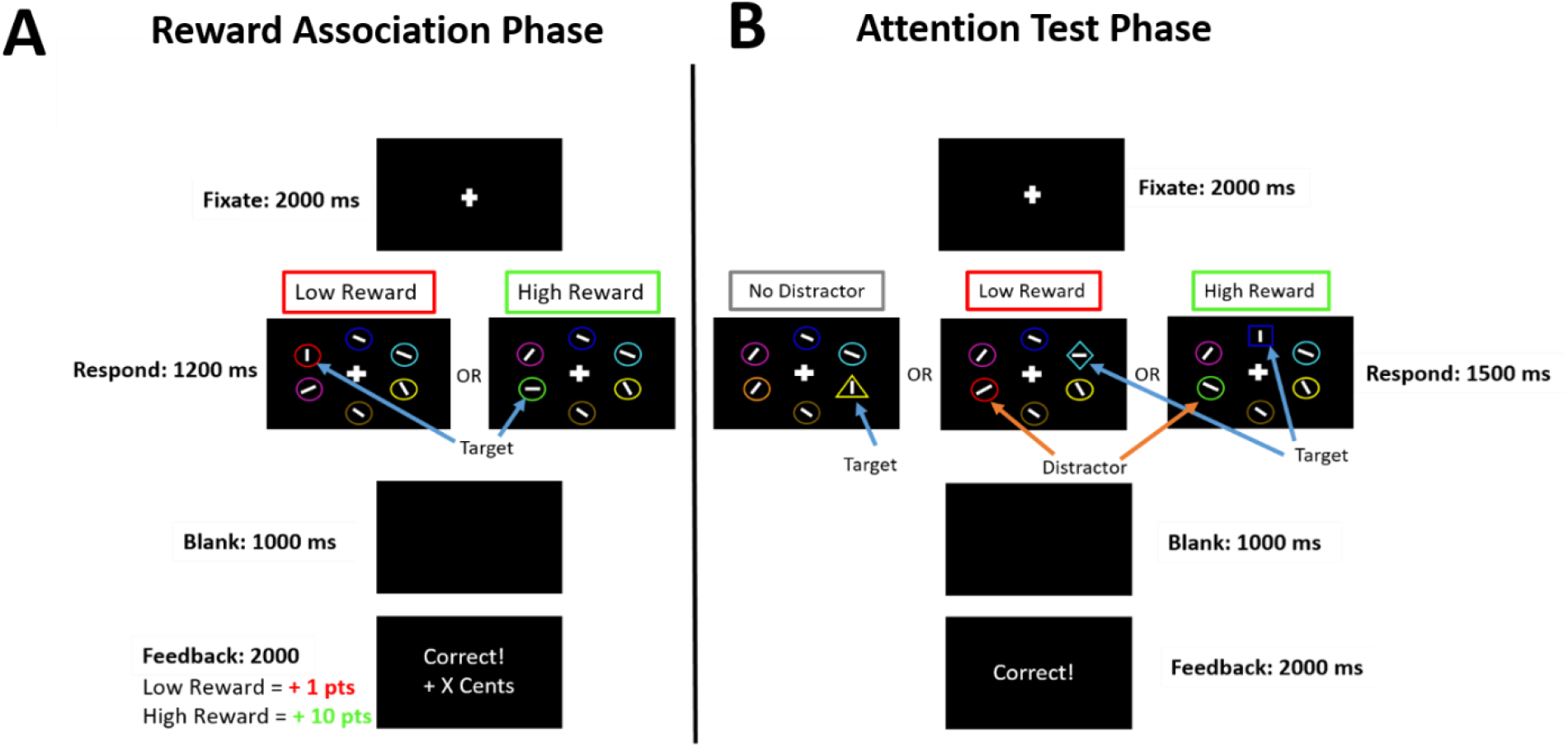
Trial sequence for the two phases of the task. (**A**) Reward association phase: Targets in the reward association phase were defined by a pre-specified color (e.g., red and green). Subjects reported the horizontal or vertical orientation of the white bar inside the target by pressing one of two keys. Correct answers were differentially rewarded based on the color (e.g., 10 cents for green and 1 cent for red). (**B**) Attention test phase: Here subjects were told to ignore the colours. Targets were identified as the unique shape. Once again, subjects reported the orientation of a white bar inside the target by pressing one of two keys. Half of the trials contained a distractor, i.e. a non-target shape in a colour previously associated with a reward (e.g., green for the low reward distractor and red for the high reward distractor). The other half of the trials did not include a distractor.

#### Reward association phase

during this phase, subjects gradually learned to associate different colors with different levels of reward. Subjects were instructed to search for a red or green target circle on a screen with five other non-target circles (2.3° x 2.3° visual angle) present surrounding a white fixation cross (0.5° x 0.5° visual angle). The stimuli were arranged in a large circle around the center of the screen with an approximate radius of 5°. Subjects were asked to report the orientation of a white bar that was inside the target circle as either vertical or horizontal using a key press. They were told to press the “z” key if the bar was vertical and the “m” key if the bar was horizontal. The bars inside the targets were always vertical or horizontal while the bars in the non-target circles were tilted at 45° to the left or right. Every trial had *either* a red or a green circle. Since subjects participated in two sessions, to avoid any carry-over effects, we used blue and yellow for the target colours in the other session. Colour assignment was counterbalanced across sessions.

At each session, one colour was assigned the high reward and one was assigned the low reward, and colour-reward assignment was counterbalanced. The high reward colour (e.g. green in **Figure 1a**) yielded 10 cents on 80% of the trials and only 1 cent on 20% of the trials upon a correct response. The low reward colour (e.g. red) led to a reward of only 1 cent on 80% of the trials and 10 cents on 20% of the trials. Feedback following incorrect responses consisted of “Incorrect!” printed in red text and “0 cents” earned. Subjects were told that they would receive a cash bonus equivalent to the winnings from 100 randomly selected trials. Subjects had 1200 milliseconds to make a response and were asked to fixate on a white cross in the center of the screen for 2000 milliseconds between each trial. If they did not respond before the time limit, they heard a loud beep and the text “Too slow!” was presented in white on the screen. Subjects performed 240 trials following 20 practice trials, and they were given the possibility of taking a break after 120 trials.

#### Attention test phase

To probe the influence of reward on attention, the previously rewarded colors (e.g. green and red in **Figure 1b**) were used as distractors in the attention test. Subjects were explicitly told that colours no longer mattered and that they should instead focus on the shape of the stimuli. In this phase, subjects were required to report the orientation (horizontal or vertical) of a white bar in a target shape. They pressed the “z’ key if the bar was vertical and “m” if the bar was horizontal. The target shape was identified as the unique shape, e.g., the diamond among circles or the circle among squares. On every trial, one target shape and five non-targets were arranged in a circle around the center of the screen. Non-target shapes also contained white bars but these were diagonally oriented at 45°. Subjects were notified if they were “Correct” or “Incorrect” with white text after making a response. Subjects had 1500 milliseconds to make a key response and, if they failed to make a response, they saw “Too Slow!” in white text accompanied by an audible beep from the computer. They were asked to fixate on a white cross in the center of the screen for 2000 milliseconds between each trial. Critically, there were three types of trials: trials with a high-reward distractor among the non-target shapes, trials with a low-reward distractor, and trials where no distractor was present. On high reward trials, the colour of one of the non-target shapes corresponded to the high-reward colour from the previous reward association phase the subject had just completed. On low reward trials, the colour of one of the non-target shapes corresponded to the low reward colour from the reward association phase. Finally, on trials where no distractor was present, none of the non-target colours corresponded to the rewarded colours. Following 10 practice trials there were 240 trials: 50% of trials included a distractor (25% were high reward, 25% were low reward), and the other 50% did not include a distractor. The order of these trials was randomized. Subjects were offered a break after 120 trials.

### Analysis

To compare performance across groups and conditions, statistics were computed using mixed effects linear and logistics regressions (R lme4 package; (Bates et al., 2015)), performed in R version 3.6.3 (R Core Team, 2020). The general approach to model specification was as follows: we included random intercepts for subjects and random slopes for all within-subject variables and interactions (e.g., medication*reward_condition) (Barr et al., 2013; Meteyard & Davies, 2020). Because the maximally specified models failed to converge, we specified a variance components structure for the G matrix that assumes zero covariance between random effects and we removed random effects that prevented a given model from converging.

For the reward association phase, to compare performance between reward conditions and groups, we performed logistic regressions with probability of a correct response on each trial as the dependent variable. We ran three separate models: one in controls only, which included only reward level (low or high) as our main experimental variable; one in Parkinson’s patients which included reward level, medication state (OFF of ON) and their interaction (medication*reward); and one in all subjects that included reward level, disease (control or Parkinson) and their interaction (disease*reward). All models included session (first or second), and Symbol Digit Modalities Test performance as covariates to control for learning and cognitive ability. The model with both patients and controls also included education as a covariate to account for sample differences. We removed sex from the reward association phase models because all attempts to include it caused convergence failures.

Analyses for the attention test phase followed a similar approach except that reaction time was the main dependent variable, which, for the sake of normality, was transformed using a base-ten logarithm. We ran three separate models: one in controls only, which included only distractor type (no distractor, low reward or high reward) as our main experimental variable; one in Parkinson’s patients which included distractor type, medication state (OFF or ON) and their interaction; and one in all subjects that included distractor type, disease (control or Parkinson) and their interaction. As above, all models included session (first or second), and Symbol Digit Modalities Test performance, and the model with both patients and controls also included education and sex as covariates to account for sample differences.

The categorical variable distractor type had three levels: high reward, low reward, and no distractor. Because we were principally interested in the differences *between* these levels, it was coded using two vectors, each with 3 levels: V1 (1=low reward, 0=high reward, −1=no distractor) and V2 (0=low reward, 1 = high reward, −1 = no distractor). As a result, the regression coefficient for V1 represented the difference in logRT between the low reward condition and the grand mean, and the regression coefficient for V2 represented the difference between the high reward condition and the grand mean. In order to test all three possible contrasts between the distractor levels (no distractor vs. low reward, no distractor vs. high reward, and low reward vs. high reward), we used the esticon function in R (Hojsgaard, 2007) to compute weighted sums of the relevant coefficients as follows: no distractor vs. low reward = 2*βV1 + βV2; no distractor vs. high reward = βV1 + 2*βV2; low reward vs. high reward = βV1 − βV2. We applied the same approach to test the contrasts between the condition*variable interactions.

We conducted follow-up analyses to evaluate overall distractibility where the three-level distractor type variable was collapsed into a new variable with only two levels (distractor present vs. absent). As above, we ran three separate models: one including only controls only, one including only patients, and one including both patients and controls.

### Hardware and Software

All computerized tasks were conducted on a MacBook Air (13-inch, 2017) with a 13.3-inch screen (diagonal) a 1440 x 900-pixel resolution and a 60Hz refresh rate. Responses were collected with the device’s built-in keyboard. Subjects sat approximately 50 cm from the display though they were instructed to take a comfortable position. Our behavioral task was coded in Python Version 2.7.

## Results

### Reward association phase

Performance in the reward association phase is presented in **Figure 2** and model estimates are presented in **Supplementary Table 2**. As expected, Parkinson’s patients ON, OFF, and controls performed well. Reward magnitude of the feedback associated with the two target colours did not affect accuracy in neither the controls (β_HC_ = 0.006, p=0.907) nor the patients (β_PD_ = 0.019, p= 0.553). Controls generally performed better than patients (β_HCvsPD_ = −0.289, p < 0.001), but the influence of reward on performance was not different between the two groups (β_disease*reward_ = 0.029, p = 0.136). Dopamine medications did not influence performance (β_ONvsOFF_ = 0.005, p = 0.938) nor did they alter the influence of reward magnitude on performance (β_med*reward_ = 0.021, p = 0.368). The absence of a difference between groups for the effect of reward on performance is important for the interpretation of the second phase of the task as it indicates that patients (in both medication conditions) and controls had similar experiences during the reward training.

**Figure 2.**
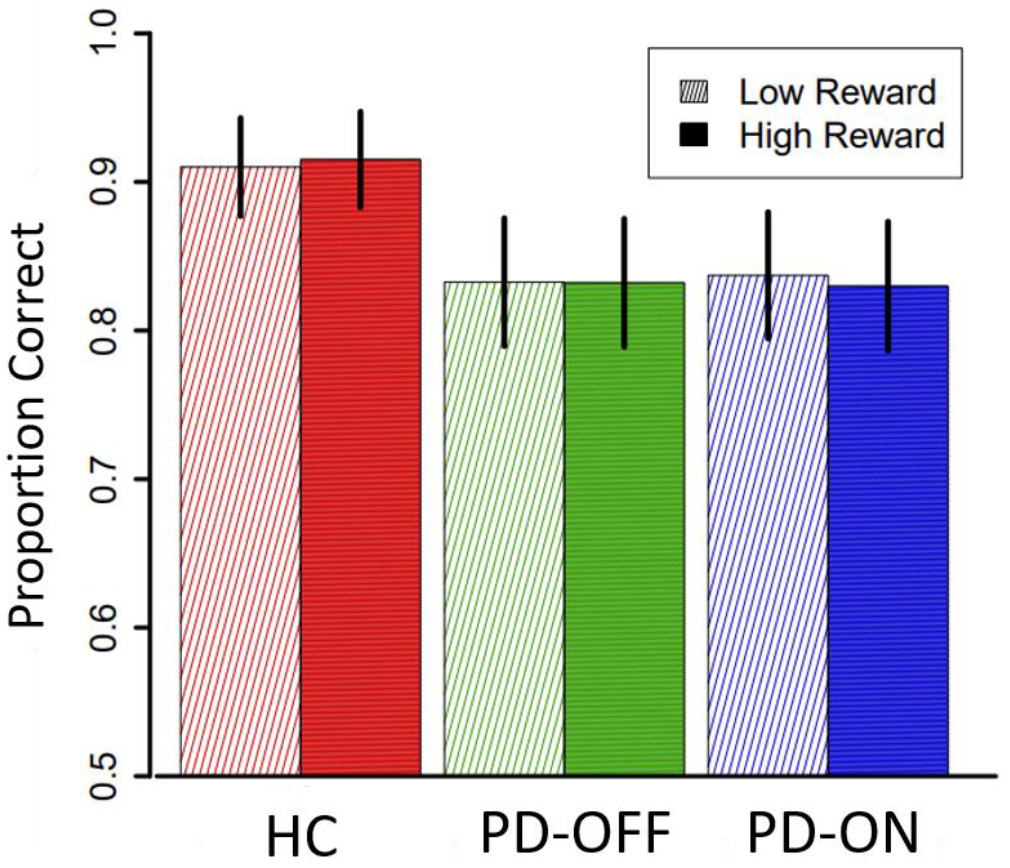
Performance during the initial colour-reward association phase. Accuracy on the initial learning phase of the task, shown separately for trials where the target colour was associated with a low reward upon correct responses versus trials where the target colour was associated with high reward upon a correct response. Controls performed better than Parkinson’s patients (p < 0.001) but importantly, there was no effect of reward level on accuracy (HC: p = 0.907; PD: p = 0.553), nor was there a difference between groups in the effect of reward magnitude on performance (HC vs. PD: p = 0.136; ON vs OFF: p = 0.368). There was no effect of dopamine state on overall performance (p = 0.938). Error bars represent the standard error of the mean.

### Attention test phase

First, we examined accuracy, which was comparable between patient drug states groups (W = 702, p = 0.411). In keeping with previous studies of the effect of reward on attention, the main outcome of interest was reaction times. Results from the attention test phase are presented in **Figure 3**, mean reaction times are presented in **Supplementary Table 3** and model estimates are presented in **Supplementary Table 4 and 5**. We were principally interested in measuring the effects of reward-associated distractors and of dopaminergic medications on attention in Parkinson’s patients. First, we found a main effect of distractors: Parkinson’s patients were slowed by the presence of both low and high reward distractors (low reward versus no distractor difference estimate = 0.010, p < 0.001; high reward versus no distractor difference estimate = 0.006, p = 0.006). However, there was no difference in slowing between low and high reward distractor trials (low vs. high difference estimate = 0.004, p = 0.170). Next, we examined whether the interaction between medication state and reward level of the distractor. Surprisingly, dopamine medications did not influence the effect of reward on attention in patients. Specifically, there was no difference between ON and OFF patients in the effect of low versus high reward distractors on reaction time (difference estimate= −0.002, p = 0.546), low reward versus no distractor (difference estimate = 0.001, p = 0.757), or high reward versus no distractor (difference estimate = 0.002, p = 0.313). The only difference between patients ON and OFF was that patients ON were slower, across all three trial types, than patients OFF (β_ONvsOFF_ = 0.008, p = 0.006). This suggests that while there was an effect of medication on response time, it was not selective to distraction.

**Figure 3.**
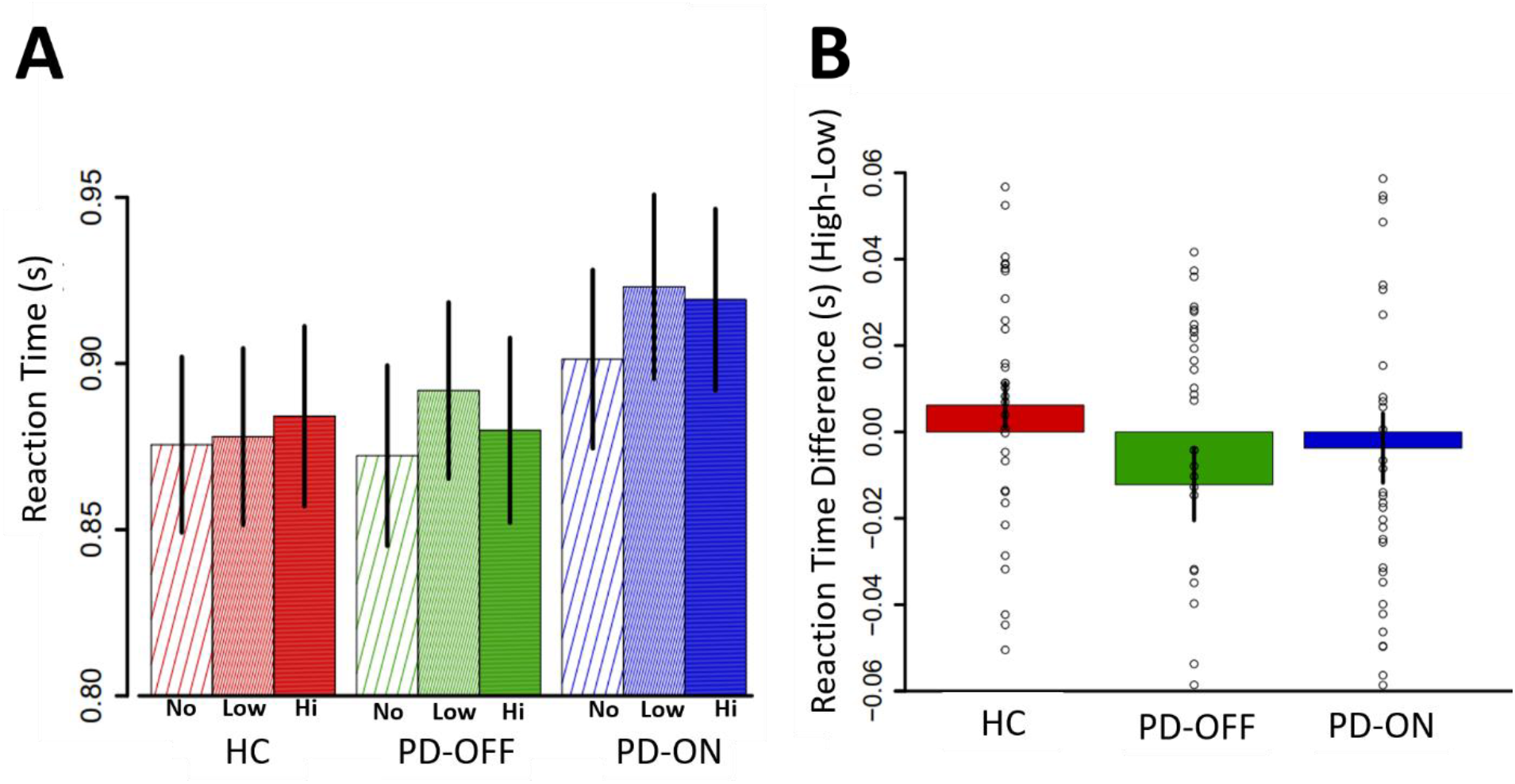
No influence of reward or dopamine on selective attention in Parkinson’s disease. (**A**) Reaction time (in seconds) on the attention task is shown for all three trial types (trials where no distractor was present, trials with a low-reward distractor and trials with a high-reward-associated distractor) for healthy controls and for patients ON and OFF medications. Parkinson’s patients were similarly slowed (i.e., distracted) by the presence of both low and high-reward distractors (low: p < 0.001, high: p = 0.006) and the degree of distraction caused by reward was not influenced by dopamine. (**B**) Difference in reaction time between the high-reward condition and the low-reward condition is shown for healthy controls and for patients ON and OFF medications. Error bars represent standard error of the mean (SEM).

To better understand these results, we also examined the effect of reward on attention within each group separately. In healthy controls, we found that though the effect of reward level on attention did not reach statistical significance (difference estimates: low vs no distractor = 0.001, p = 0.530; high vs. none = 0.004, p = 0.100, low vs. high = −0.002, p = 0.376), the overall direction of the effect was as expected: controls were slowest on the trials that included a high-reward distractor (881ms ± 235), compared to the trials with a low-reward distractor (875ms ± 233) and those without a distractor (871ms ± 230). Furthermore, the degree of slowing induced by the highly-rewarded distractor was comparable to that reported previously in young controls (Anderson, 2013). In contrast, Parkinson’s patients OFF were significantly slowed only by the low-reward distractor (low vs. none difference estimate = 0.009, p=0.005) and patients ON were slowed by both the low-and high-reward distractors, but to a similar degree (difference estimates: low vs. none = 0.01, p=0.001; high vs. none = 0.008, p=0.008; low vs. high = 0.002, p=0.587).

In order to measure effects of disease on performance, we also compared patients to controls. Overall, patients were not slower than controls (β_HCvsPD_ = 0.001, p = 0.863). With respect to the effects of interest, Parkinson’s patients were more distracted than controls by the presence of a low reward distractor (low vs. no distractor difference estimate = 0.004, p = 0.008). However, this pattern was not observed in the presence of a high reward distractor (high vs. no distractor difference estimate = 0.001, p = 0.423). As noted above, all models controlled for session and performance on the Symbol Digit Modalities Test. Overall, across both groups, participants were faster on the second session (β_session_ = −0.012, p<0.001), and better performance on Symbol Digit Modality Test was associated with faster responses (β_SDMT_ = −0.002, p<0.001).

We were also interested in the effect of disease and dopaminergic medications on overall distractibility. To examine this, we collapsed across reward levels and compared reaction times on trials with a distractor to reaction times on trials without a distractor (**Supplementary Table 6.**). We found that patients were more distractible than controls (β_disease*distraction_ = 0.001, p = 0.035), however, we did not find an effect of dopaminergic medication on distractibility (β_drugs*distraction_ = 0.001, p = 0.417).

Finally, due to known links between mood symptoms and reward sensitivity deficits in Parkinson’s disease (Muhammed et al., 2016), we looked at the relationship between apathy and depression symptoms (measured with the Apathy Evaluation Scale and the Geriatric Depression Scale respectively) and the extent to which reward caused distraction across patients (**Supplementary Figure 1.**). We did not find any significant relationship between apathy and reward-driven attention (ρ_OFF_=0.15, p = 0.385; ρ_ON_=-0.28, p = 0.077), nor between depression and reward-driven attention (ρ_OFF_=-0.09, p = 0.598; ρ_ON_=0.12, p = 0.463).

## Discussion

Reward is known to exert an automatic, involuntary effect on the allocation of attentional resources (Anderson, 2013; Anderson et al., 2017). In Parkinson’s disease, both reward signaling deficits (Bódi et al., 2009; Frank et al., 2004; Muhammed et al., 2016) and attention impairments are well established (Dirnberger & Jahanshahi, 2013; Robbins & Cools, 2014). However, little is known about whether the reward deficits in patients directly contribute to poor attention, and more specifically, whether Parkinson’s patients suffer from a dopamine-dependent inability to use reward to selectively allocate their attention resources. Using a task where the presence of reward-associated distractors was used to probe the automatic reward-driven allocation of attention, we found that Parkinson’s patients were insensitive to reward information during the allocation of their attention. High-reward stimuli did not lead to greater attention capture than low reward stimuli in patients. However, contrary to our predictions, we did not find evidence that dopamine modulated this inability to use reward to guide attention allocation. By showing an inability to selectively allocate attention based on reward in Parkinson’s patients, our results provide a novel mechanism by which attention might be impaired in Parkinson’s disease.

Our findings are broadly consistent with the body of work showing that patients are impaired at goal-directed attention allocation (Fallon et al., 2016; Owen et al., 1993; Rustamov et al., 2014; Sawada et al., 2012; Slabosz et al., 2006) and extend these findings by showing that even the more automatic prioritization of attention by reward is impaired in Parkinson’s patients. Goal-directed, or top-down control of attention has typically been studied in Parkinson’s patients by measuring attentional set formation (Fallon et al., 2013; Fallon et al., 2016; Owen et al., 1993b; Slabosz et al., 2006). In the typical task used, participants are explicitly aware of which stimuli they must attend to (the attentional set) and successful performance depends on the ability to adequately form, maintain, and switch attentional sets according to those explicit task goals. The current study focused instead on the more automatic, or involuntary, mechanism by which attention is allocated to behaviourally-relevant stimuli. Such a mechanism is essential for matching the available cognitive resources to the nearly infinite amount of information constantly generated by our environment, not all of which is goal-relevant at a given time. In order to disentangle reward-guided attention from top-down goal-directed modulation, we used a task where reward was exclusively paired with distractors, and was therefore in direct opposition to task goals. Previous studies using this task have found a relationship between striatal function and reward-guided attention selectivity in healthy individuals (Anderson, 2017; Anderson et al., 2014), suggesting that loss of normal striatal reward signaling in Parkinson’s patients may underlie the loss of reward-guided attention selectivity.

Though a relationship between striatal dysfunction and an inability to use reward to guide attention allocation seems highly plausible in Parkinson’s patients, the exact mechanism linking the two is less clear. One possibility is that, given known reinforcement learning deficits in Parkinson’s disease (Bódi et al., 2009; Cools et al., 2006; Frank et al., 2004; Palminteri et al., 2009; Rutledge et al., 2009; Skvortsova et al., 2017), patients do not learn the reward values of the color-stimuli in the first place. This could then lead to a blunting of the differential levels of attention capture caused by the colours. While it seems likely that reward learning deficits could generally impact selective attention in Parkinson’s patients, we found no evidence in the current task that this was the case. First, to ensure that learning deficits would not interfere, the learning phase was designed to be easy: average accuracy in the patients was 85.8%. Second, though the task did not allow us to specifically assess the strength of the colour-reward associations formed in the first phase, accuracy on the high reward and low reward trials were equivalent in all groups, indicating that learning performance was not biased by the different levels of reward. This suggests that the selective attention impairment found in the present study cannot be explained solely on the basis of reduced initial responsiveness to reward or inability to learn from it. Another possibility is that the impairment occurs at the stage of perceptual processing of the stimuli. Indeed, reward is thought to modulate attention by facilitating the perceptual processing of reward-associated stimuli, an effect that appears to depend on striatal activation *while* the selective allocation of attention is occurring, i.e. after the initial learning of the reward value has occurred, and at a time when the reward information is no longer relevant to the task (Hickey & Peelen, 2015). Though exactly how this enhanced (or in the case of patients, possibly blunted) perceptual processing occurs is not clear, it has been proposed that the striatum plays a role in gating of information through the balance of activation in direct and indirect pathways (Chatham & Badre, 2015; Frank et al., 2001; Frank & O’Reilly, 2006; Hazy et al., 2007). This gating mechanism has mainly been suggested to explain the flow of information into and out of working memory, but could also have implications for the automatic selection of information to be processed in attention in Parkinson’s disease (Frank, 2005). One possibility is that dopaminergic dysregulation produces a deficit in reward-oriented gating of attention such that rewarding signals which normally “open” the gate no longer have such an effect. This could explain why patients do not allocate their attention more towards high reward than low reward stimuli. However, potentially complicating the above interpretation that the striatum is involved in reward-guided attention allocation is the fact that we did not find an effect of dopamine replacement on reward-guided attention in patients.

Our finding that reward-guided attentional selectivity was not sensitive to dopamine replacement was unexpected. We think this should be interpreted in the specific context of Parkinson’s disease, where the effects of dopamine on cognition are quite complex, and does not necessarily generalize to the role of dopamine in healthy brain function. One possible explanation is that top-down, goal-directed attention deficits, and in particular the ability to *shift* attention, which is known to be impaired in Parkinson’s patients (Fallon et al., 2016; Slabosz et al., 2006), might have made it more difficult to detect the more subtle effects of reward on attention allocation. Attentional shifting deficits are present both in the OFF and ON medication state: patients OFF tend to perseverate on previously relevant stimuli (Fallon et al., 2013), whereas patients ON tend to have difficulty shifting their attention towards previously irrelevant features (Owen et al., 1993a). Inherent to our task was the requirement to explicitly shift attentional focus: from colours during the learning phase to shapes during the test phase. It is therefore possible that a tendency in the patients OFF to perseverate on the now irrelevant colours and an inability in the patients ON to shift their attention to the previously irrelevant shapes impacted performance during the test phase, and that the presence of these top-down attentional control deficits overrode the more subtle dopamine-dependent effects of reward on automatic attention allocation. Thus, it is possible that both reward-driven selectivity and top-down attentional set formation are impaired in Parkinson’s patients. Another possible explanation is that dysregulation of other neuromodulator systems contributes to selective attention deficits in Parkinson’s patients, and possibly even interacts with the deficits caused by dopaminergic loss, making it harder for dopamine replacement alone to remediate impairments. For instance, loss of cortical cholinergic projections from the basal forebrain is known to occur early in the disease and has been associated with attentional deficits (Bohnen et al., 2006; Lee et al., 2019). In particular the ability to resist distraction appears to be dependent on cholinergic integrity more so than dopaminergic integrity (Kim et al., 2017). Consistent with this, we found that dopamine state did not influence attention capacity, measured here as overall distractibility. It will be important for future studies to use tasks where resistance to distraction (i.e., top-down attentional control) is not in direct opposition to reward-driven attentional selectivity in order to obtain more sensitive measurements of automatic attention allocation, and to consider pharmacological manipulations of other neurotransmitter systems to attempt to triangulate the specific contributions of each.

It is important to note that in our sample of older controls, we did not find the clear effect of reward magnitude on attention selectivity that has been repeatedly shown in younger controls (Anderson, 2013; Anderson et al., 2011; Anderson & Halpern, 2017; Anderson & Yantis, 2013). Though the difference in the slowing of responses between the high reward and low reward conditions was not statistically significant, it is worth noting that this difference (6 ms) was of similar magnitude to that reported previously in young controls (Anderson, 2013; Anderson et al., 2011; Anderson & Halpern, 2017; Anderson & Yantis, 2013). However, as is typical of older adult samples, the variability was considerably higher, suggesting that we were underpowered to detect the desired effects. A weaker and more variable effect of reward could arise from two possible sources: an age-related reduction in sensitivity to rewarding outcomes (Eppinger et al., 2011), possibly related to a decline in midbrain dopaminergic function (Kaasinen & Rinne, 2002), and age-related differences in the power of monetary rewards to act as incentives (Rademacher et al., 2014; Samanez-Larkin et al., 2007; Spaniol et al., 2015), which we used to drive reward-colour associations as has been done previously. Given that the mean age of both our patient and control samples was over 60, we cannot discount the possibility of age-related blunting of reward sensitivity. To our knowledge, reward-driven attention has not previously been tested in aging populations so future work might shed light on this question.

An important limitation to our study is that the task we used, and others like it, tend to elicit relatively small effect sizes (Anderson, 2013; Anderson et al., 2011; Anderson & Halpern, 2017; Anderson & Yantis, 2013; Gong & Li, 2014; Klink et al., 2017), especially when compared to those related to top-down control of attention, detailed above. In our case, as in previous reports of this task, changes in reaction time were on the order of ten or twenty milliseconds. Effect sizes of this magnitude are difficult to detect in populations whose cognitive behavior is inherently noisy, such as older adults and neurological patient populations in particular (Burton et al., 2006; Hultsch et al., 2002). Future work could incorporate peripheral physiological measures such as eye-tracking to improve sensitivity of the outcome measures.

In summary, we found that Parkinson’s patients did not effectively use reward to guide the allocation of their attention, but, contrary to predictions, that this impairment was not modulated by dopamine state. This extends previous attention work by showing that in addition to impairments in the voluntary allocation of attention, Parkinson’s patients are impaired in the automatic allocation of attention to behaviourally relevant stimuli in the environment. A stimulus’ ability to capture attention marks its priority for downstream cognitive processing. The voluntary and involuntary mechanisms that support the ability to selectively attend to behaviourally relevant information can therefore reasonably be viewed as basic elements of adaptive behaviour. An important future direction will therefore be to examine the relationship between impaired selective attention and other core cognitive deficits of Parkinson’s disease – impaired working memory, learning, and decision-making – in order to determine whether they share a common source. The identification of such a common source would inform the development of cognitive rehabilitation approaches that could have broad impact.

## Acknowledgments

We thank Léah Suissa-Rocheleau and Elsie Yan for help with participant testing and we thank the participants for their time and interest.

## Supplementary Material

**Supplementary Table 1.** Demographic and clinical characteristics of Parkinson’s patients ON and OFF medication. This information is presented in addition to Table 1 in the main text because most, but not all patients, completed both the ON and OFF sessions. The groups presented below reflect the groups included in the analyses comparing patients ON vs. OFF. Twenty-eight of these patients were tested in both conditions.

| Variable | Patients ON (N = 40) | Patients OFF (N = 35) | P-Value |
| --- | --- | --- | --- |
| Age | 63.5 (6.4) | 64.2 (5.9) | .636 |
| Education, years | 15.6 (3.2) | 15.4 (3.3) | .813 |
| Disease duration, years | 4.8 (3.4) | 5.1 (3.4) | .738 |
| Total PD Medication, mg | 674.2 (336.7) | 667.0 (332.0) | .926 |
| % Taking Dopamine Agonists | 14% | 17% | .999 |
| % Female | 23% | 33% | .503 |
| MoCA | 27.1 (2.2) | 27.0 (2.5) | .927 |
| Verbal Fluency (MoCA) | 12.4 (4.3) | 12.6 (4.0) | .873 |
| Digit Span Test | 11.2 (2.3) | 11.2 (2.1) | .961 |
| Symbol Digit Modalities Test | 40.9 (10.5) | 40.7 (10.5) | .923 |
| Geriatric Depression Scale | 7.9 (6.1) | 7.6 (5.8) | .815 |
| Apathy Evaluation Scale | 58.5 (8.01) | 58.6 (7.9) | .945 |

MoCA = Montreal Neurological Assessment, Verbal Fluency is taken from the Language section of the MoCA. Values presented are mean (SD).

**Supplementary Table 2.** Reward Association phase model coefficient estimates. Fixed effects estimates obtained from the mixed effects regression indicating the influence of trial type, medication or disease, and other covariates on response accuracy during the Reward association phase. Coefficients are presented separately for the three models that were run: PD Only to measure the effect of medications, HC Only, and All Subs to measure the difference between patients and controls.

| Fixed effect | Model | Estimate (SE) | P-Value |
| --- | --- | --- | --- |
| Intercept | PD Only | 0.038 (0.370) | 0.918 |
| Medication | PD Only | 0.005 (0.063) | 0.938 |
| Reward Level | PD Only | 0.019 (0.031) | 0.553 |
| Medications * Reward Level | PD Only | 0.021 (0.024) | 0.368 |
| Session | PD Only | 0.042 (0.064) | 0.513 |
| SDMT | PD Only | 0.042 (0.009) | <0.001 *** |
| Intercept | HC Only | 1.518 (0.824) | 0.056 |
| Reward Level | HC Only | 0.006 (0.053) | 0.907 |
| Session | HC Only | 0.091 (0.090) | 0.312 |
| SDMT | HC Only | 0.022 (0.017) | 0.190 |
| Intercept | All Subs | 0.073 (0.501) | 0.885 |
| Disease | All Subs | -0.289 (0.087) | <0.001 *** |
| Reward Level | All Subs | -0.004 (0.020) | 0.8256 |
| Disease * Reward Level | All Subs | 0.029 (0.020) | 0.136 |
| Session | All Subs | 0.054 (0.0529) | 0.308 |
| SDMT | All Subs | 0.032 (0.008) | <0.001 *** |
| Education | All Subs | 0.041 (0.027) | 0.121 |

SDMT = Symbol Digit Modalities Test

**Supplementary Table 3.** Mean reaction times during the Attention test phase. Mean reactions times (in seconds) are presented for the three trial types (no distractor, low reward and high reward) for the three groups. In the case of the Control group reaction times are averaged across both sessions. Vales are mean (SD).

| Group | Medication Status | Reward Level | Reaction Time (s) |
| --- | --- | --- | --- |
| Control | NA | None | 0.871 (0.230) |
| Control | NA | Low | 0.875 (0.233) |
| Control | NA | High | 0.881 (0.235) |
| Parkinson's | OFF | None | 0.894 (0.237) |
| Parkinson's | OFF | Low | 0.907 (0.237) |
| Parkinson's | OFF | High | 0.895 (0.235) |
| Parkinson's | ON | None | 0.905 (0.236) |
| Parkinson's | ON | Low | 0.916 (0.237) |
| Parkinson's | ON | High | 0.908 (0.236) |

**Supplementary Table 4.** Attention Test phase - model coefficient estimates. Fixed effects estimates obtained from the mixed effects regression indicating the influence of trial type, medication or disease, and other covariates on the reaction times (log(RT)) during the Attention Test phase. Coefficients are presented separately for the three models that were run: PD Only to measure the effect of medications, HC Only, and All Subs to measure the difference between patients and controls.

| Fixed effect | Model | Estimate (SE) | P-Value |
| --- | --- | --- | --- |
| Intercept | PD Only | 0.033 (0.026) | 0.224 |
| Reward Vector 1 | PD Only | 0.004 (0.001) | 0.002 ** |
| Reward Vector 2 | PD Only | 0.001 (0.014) | 0.553 |
| Medication | PD Only | 0.008 (0.003) | 0.006 ** |
| Reward Vector 1* Medication | PD Only | -0.000 (0.001) | 0.840 |
| Reward Vector 2* Medication | PD Only | 0.001 (0.001) | 0.373 |
| Session | PD Only | -0.006 (0.003) | 0.021 * |
| SDMT | PD Only | -0.002 (0.001) | <0.001 *** |
| Intercept | HC Only | 0.014 (0.047) | 0.772 |
| Reward Vector 1 | HC Only | -0.000 (0.001) | 0.838 |
| Reward Vector 2 | HC Only | 0.002 (0.001) | 0.163 |
| Session | HC Only | -0.016 (0.002) | <0.001 *** |
| SDMT | HC Only | -0.002 (0.001) | 0.089 |
| Intercept | All Subs | 0.010 (0.033) | 0.751 |
| Reward Vector 1 | All Subs | 0.002 (0.001) | 0.041 * |
| Reward Vector 2 | All Subs | 0.001 (0.001) | 0.163 |
| Disease | All Subs | 0.001 (0.006) | 0.863 |
| Reward Vector 1 * Disease | All Subs | 0.002 (0.001) | 0.019 * |
| Reward Vector 2 * Disease | All Subs | -0.001 (0.001) | 0.585 |
| Session | All Subs | -0.012 (0.002) | <0.001 *** |
| SDMT | All Subs | -0.002 (0.001) | <0.001 *** |
| Education | All Subs | 0.002 (0.002) | 0.193 |
| **Sex** | All Subs | -0.017 (0.011) | 0.137 |

SDMT = Symbol Digit Modalities Test

**Supplementary Table 5.**
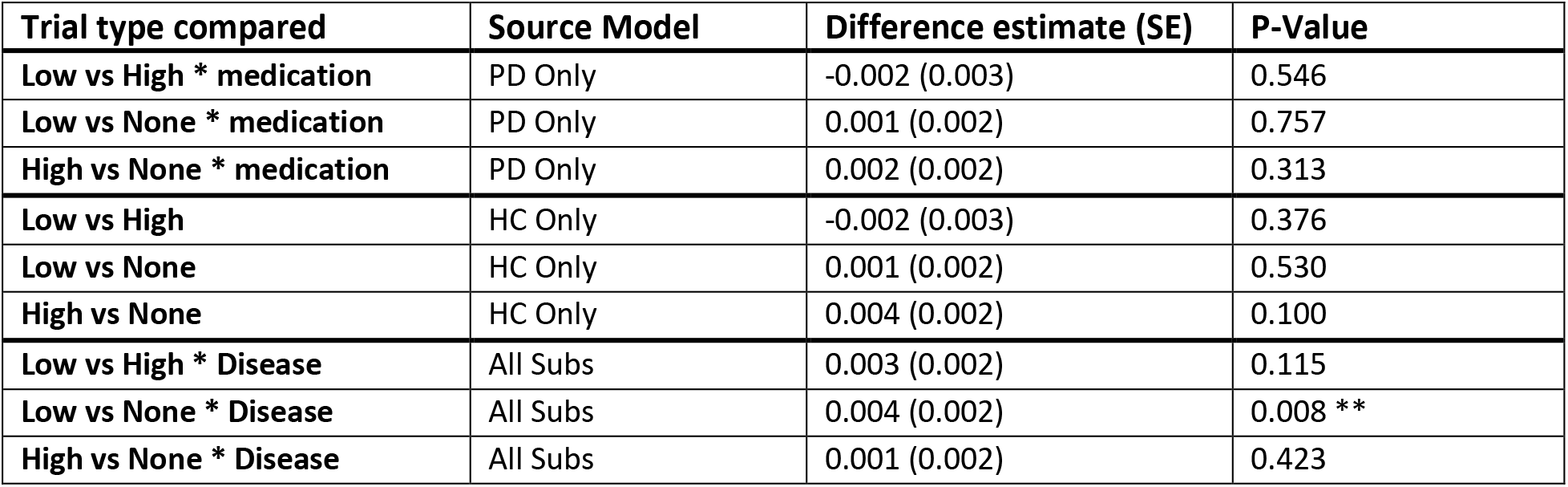
Attention test phase analysis to look at effect of reward level and disease/medication. Difference estimates computed from weighted sums of the regression coefficients presented in Supplementary Table 4 in order to obtain the effect of trial type on reaction times, and the difference between groups for this effect.

| **Trial type compared** | **Source Model** | **Difference estimate (SE)** | **P-Value** |
| --- | --- | --- | --- |
| **Low vs High \* medication** | PD Only | -0.002 (0.003) | 0.546 |
| **Low vs None \* medication** | PD Only | 0.001 (0.002) | 0.757 |
| **High vs None \* medication** | PD Only | 0.002 (0.002) | 0.313 |
| **Low vs High** | HC Only | -0.002 (0.003) | 0.376 |
| **Low vs None** | HC Only | 0.001 (0.002) | 0.530 |
| **High vs None** | HC Only | 0.004 (0.002) | 0.100 |
| **Low vs High \* Disease** | All Subs | 0.003 (0.002) | 0.115 |
| **Low vs None \* Disease** | All Subs | 0.004 (0.002) | 0.008 ** |
| **High vs None \* Disease** | All Subs | 0.001 (0.002) | 0.423 |

**Supplementary Table 6.** Attention test phase analysis to look at effect of overall distraction – model estimates. Fixed effects estimates obtained from the mixed effects regression indicating the influence of trial type, medication or disease, and other covariates on the reaction times (log(RT)) during the Attention Test phase. Coefficients are presented separately for the three models that were run: PD Only to measure the effect of medications, HC Only, and All Subs to measure the difference between patients and controls.

| **Fixed effect** | **Model** | **Estimate (SE)** | **P-Value** |
| --- | --- | --- | --- |
| **Intercept** | PD Only | 0.031 (0.026) | 0.241 |
| **Medication** | PD Only | 0.007 (0.003) | 0.007 ** |
| **Distractor Presence** | PD Only | 0.004 (0.001) | <0.001 *** |
| **Medication \* Distractor Presence** | PD Only | 0.001 (0.001) | 0.421 |
| **Session** | PD Only | -0.006 (0.003) | 0.002 * |
| **SDMT** | PD Only | -0.002 (0.001) | <0.001 *** |
| **Intercept** | All Subs | 0.010 (0.033) | 0.767 |
| **Disease** | All Subs | 0.001 (0.006) | 0.923 |
| **Distractor Presence** | All Subs | 0.003 (0.001) | <0.001 *** |
| **Disease \* Distractor Presence** | All Subs | 0.001 (0.001) | 0.035 |
| **Session** | All Subs | -0.012 (0.002) | <0.001 *** |
| **SDMT** | All Subs | -0.002 (0.001) | <0.001 *** |
| **Education** | All Subs | 0.002 (0.002) | 0.195 |
| **Sex** | All Subs | -0.017 (0.011) | 0.136 |

SDMT = Symbol Digit Modalities Test

**Supplementary Figure 1.**
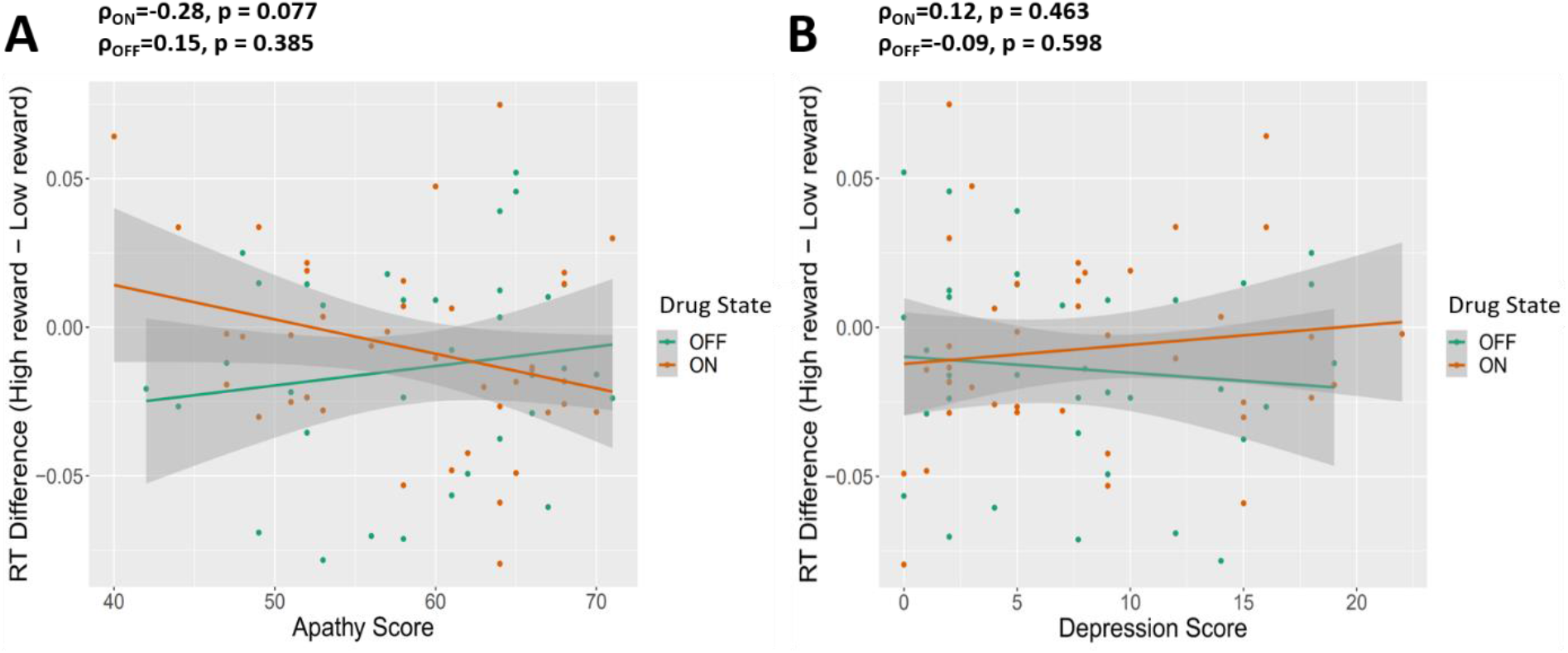
Individual differences in reward driven attention allocation are not related to apathy or depression in Parkinson’s patients. The effect of reward on attention, measured as the difference in reaction time in seconds between trials with a high reward distractor versus trials with a low reward distractor, is plotted separately for performance when tested OFF and ON dopaminergic medications, and is plotted against scores on the Apathy Evaluation Scale scores (A) and Geriatric Depression Scale (B).

